# *Magel2* in hypothalamic POMC neurons influences the impact of stress on anxiety-like behavior and spatial learning associated with a food reward in male mice

**DOI:** 10.1101/2025.07.09.663942

**Authors:** Sangbhin Lee, Young-Hwan Jo

## Abstract

Prader-Willi syndrome (PWS) results from a lack of expression in several paternally inherited, imprinted contiguous genes. Among the genes inactivated in PWS, the *Magel2* gene is considered a significant contributor to the etiology of the syndrome. The loss of the *Magel2* gene causes abnormalities in growth and fertility and increased adiposity with altered metabolism in adulthood, which aligns with some of the pathologies observed in PWS. Given that anxiety is a prominent phenotypic behavior in PWS, we investigate the role of the *Magel2* gene, particularly in hypothalamic POMC neurons innervating the medial amygdala (MeA), in the behavioral phenotypes associated with Prader-Willi Syndrome (PWS). Both male and female mice lacking the *Magel2* gene in MeA-innervating ARC^Pomc^ neurons display no alterations in anxiety-like behavior during the open field test, light/dark test, and elevated plus maze test in the absence of exposure to acute stress. However, male mice with a *Magel2* gene deletion in these particular neurons exhibit increased stress-induced anxiety-like behavior and reduce motivation/spatial learning, while female mice do not show these behavioral changes. Our results suggest that the *Magel2* gene in ARC^Pomc^ neurons, especially in males, influences the impact of stress on anxiety-like behavior and spatial learning deficits associated with a food reward. With the recent approval of a novel treatment for hyperphagia in PWS by the FDA that seems to target the hypothalamic melanocortin system, understanding the cellular mechanisms by which MAGEL2 in ARC^Pomc^ neurons innervating the MeA regulates emotional behaviors might help the development of new therapeutic strategies for addressing mental illness in individuals with PWS.

## Introduction

Prader-Willi Syndrome (PWS) is a neurogenetic disorder that results from the loss of several paternally inherited, imprinted contiguous genes in the human 15q11-q13 and mouse 7C regions [1; 2; 3; 4; 5]. One of the defining characteristics of PWS is an insatiable appetite for food, which can cause substantial weight gain in children during the early stages of development [6]. In addition to hyperphagia and obesity, individuals with PWS are also highly susceptible to mental health disorders [7]. Key phenotypic PWS behaviors include temper outbursts, anxiety, obsessive-compulsive behaviors, and social cognition deficits [7; 8]. These behaviors have a significant impact on the daily functioning and quality of life for individuals with PWS and their families [7]. Hence, it is necessary to determine which imprinted gene is responsible for these mental disorders in PWS.

Among the genes inactivated in PWS, the *Magel2* gene is highly expressed in the hypothalamus [9; 10; 11; 12]. Amongst hypothalamic neurons, proopiomelanocortin (POMC) neurons in the arcuate nucleus of the hypothalamus (ARC) play a critical role in controlling energy balance and mood disorders [13]. Intriguingly, *Magel2*-deficient mice have fewer ARC^Pomc^ neurons [14] and lower levels of α-melanocyte-stimulating hormone (α-MSH) derived from POMC [15]. ARC^Pomc^ axonal projections to the paraventricular and dorsomedial hypothalamus are significantly reduced by *Magel2* deletion [11].Furthermore, the basal spontaneous activity of ARC^Pomc^ neurons is lower in *Magel2*-null mice than in controls [16]. These prior studies support that MAGEL2 is critical for developing and maintaining ARC^Pomc^ neuronal circuits. In our prior studies [17; 18], we showed that optogenetic stimulation of the ARC^Pomc^ neurons −> the medial amygdala (MeA) pathway reduced acute food intake. More importantly, we also demonstrated that deleting the *Magel2* gene exclusively in ARC^Pomc^ neurons that project to the MeA inhibited the onset of diet-induced obesity (DIO) in both male and female mice during a high-fat diet (HFD) feeding, which seems to be linked to an increase in physical activity [18]. This finding is not consistent with the expectation that the loss of function of the PWS imprinted genes leads to the development of obesity, as observed in individuals with PWS. However, mice deficient in the paternal allele of *Magel2* did not develop DIO when maintained on HFD [19]. Hence, these prior findings suggest that the *Magel2* gene is unlikely to play a major role in regulating feeding behavior in PWS.

Acute emotional stressors, such as restraint and forced swim, trigger the expression of *c-fos* mRNA, a marker of neuronal activity, in most ARC^Pomc^ neurons as well as in the MeA [20]. The increased activity of ARC^Pomc^ neurons induces anxiety-like behavior [20]. Given that anxiety is a prominent phenotypic behavior in PWS [7], it is plausible that the *Magel2* gene in ARC^Pomc^ neurons, which project to the MeA, might be involved in the onset of anxiety rather than regulating feeding behavior in individuals with PWS. Hence, we sought to investigate the behavioral implications of the dysfunction of the *Magel2* gene in ARC^POMC^ neurons that innervate the MeA and to provide new and valuable insights into the role of central MAGEL2 in the regulation of emotional behaviors.

## Materials and Methods

### Ethics statement

All mouse care and experimental procedures were approved by the Institutional Animal Care Research Advisory Committee of the Albert Einstein College of Medicine and were performed following the guidelines described in the NIH guide for the care and use of laboratory animals. Stereotaxic surgery and viral injections were performed under isoflurane anesthesia.

### Animals

Rosa26-floxed-stop Cas9-eGFP (Rosa26-Cas9^f^, stock #026175) was purchased from the Jackson Laboratory. Mice were housed in cages at a controlled temperature (22 °C) with a constant humidity (40-60%) and a 12:12h light-dark cycle. Mice were fed a standard chow diet with *ad libitum* access to water.

### Viral injection

We employed neuronal *Pomc* enhancers 1 and 2 (nPEs) that were identified by the Rubinstein research group [21] to induce the expression of the Cre recombinase solely in ARC^Pomc^ neurons in adult mice, a method that has proven effective in our prior studies [22; 23]. Retrograde AAV-nPEs-Cre-WPRE viruses (serotype retrograde) were made at Applied Biological Materials, Inc (ABM). To knock down the *Magel2* gene in ARC^Pomc^ neurons innervating the MeA, retrograde AAV-PGK-loxp-tdTomato-loxp-U6-mouse *Magel2* sgRNA viruses were also generated at ABM. The sgRNAs target the consensus coding sequence 52264.1 region of mouse *Magel2* (NM_013779.2). The sequences of *Magel2* sgRNA were the following: (1) sgRNA1: cgcagctaagtacgaatctg, (2) sgRNA2: gtagggcggctatggactgc, and (3) sgRNA3: atggtccaggctccaccgct. The effectiveness of these sgRNAs was validated in our prior study [18].

A total of 84 mice were divided into four groups: 20 mice in the male control group, 20 mice in the male experimental group, 19 mice in the female control group, and 25 mice in the female experimental group. 6 and 7-week-old mice (males, ~ 20 g and females, ~18 g) were anesthetized deeply with 3% isoflurane and placed in a stereotaxic apparatus (David Kopf Instruments). A deep level of anesthesia was maintained throughout the surgical procedure. Under isoflurane anesthesia (2%), a total of 300 nl (150+150 nl from each) of retrograde AAV-PGK-loxp-tdTomato-loxp-U6-mouse *Magel2* sgRNA viruses (titer, 1 × 10^12^ GC/ml) along with retrograde AAV-nPE-Cre-WPRE (titer, 1.2 × 10^13^ GC/ml) were bilaterally injected into the MeA of Rosa26-Cas9^f^ mice (AP, −1.58 mm; ML, ± 2 mm; DV, −5 mm). A 2.5 ul Hamilton syringe, having a 33G needle, was used to inject a volume of 50 nl viruses every 10 min. The Hamilton syringe tip was left in place for 10 min after delivering viruses in order to prevent the backflow of the viral solution up the needle track.

We conducted a systematic examination of GFP expression as an indicator of viral transfection following behavioral experiments. To achieve this, transverse brain slices were prepared as described in our prior study [24]. The animals were aHnesthetized using isoflurane, and following decapitation, the brain was transferred into a sucrose-based solution, which was bubbled with 95% O2/5% CO2 and maintained at 3°C. This solution comprised the following components (in mM): 248 sucrose, 2 KCl, 1 MgCl_2_, 1.25 KH_2_PO_4_, 26 NaHCO_3_, and 10 glucose. Transverse coronal brain slices (200 μm) were prepared using a vibratome (Leica VT 1000S). The brain slices were then placed on the stage of an upright, infrared-differential interference contrast microscope (Olympus BX50WI) mounted on a Gibraltar X-Y table (Burleigh), and GFP-positive cells were visualized with a 40X water immersion objective using infrared microscopy (DAGE MTI camera). Data were excluded if no GFP-positive cells were observed in the sections.

### Immunostaining

Mice were anesthetized with isoflurane (3%) and transcardially perfused with 4% paraformaldehyde. Brain samples were post-fixed in 4% paraformaldehyde overnight in a cold room and then in 30% sucrose the following day. Tissues were sectioned using a cryostat at 20 μm. The sections were incubated with 0.3% Triton X-100 at room temperature (RT) for 30 min, and then blocked in PBS buffer containing 5% donkey serum, 2% bovine serum albumin and 0.15% Triton X-100 for 1hr at RT and then incubated with goat anti-GFP (NB100-1770, Novus), rabbit anti-POMC (1:1000, H-029-30, Phoenix biotech), and rabbit anti-MAGEL2 (1:2000, a gift from Dr. Tacer [15]) antibodies for overnight at a cold room, and then sections were washed three times in PBS and incubated with Alexa 488 anti-goat IgG for GFP (1:1000, 705-545-147, Jackson immunoresearch), Alexa 594 anti-rabbit IgG for POMC and MAGEL2 (1:1000, 711-585-152, Jackson immunoresearch) for 3hr at RT. After washing, the sections were stained with DAPI and mounted using VECTASHIELD medium (H-2000, Vector Lab.). Images were acquired using a Leica SP8 confocal microscope at a magnification of 20X. Using the cell count plugin in ImageJ software (version FIJI), MAGEL2-positive neurons were counted within the ARC, specifically in an area measuring 400μm by 400μm.

### Animal behavior tests

For restraint stress, mice were restrained for 30 min in a 50 ml conical tube with the holes (0.5 cm in diameter) at the front and back for air flowing. Mice were able to move their head and limb but not their body, and unable to access food and water during the restraint. After the restraint was completed, mice were performed behavior tests immediately.

#### Open field exploration test

Mice were placed in the center of a chamber measured 40 cm (length) x 40 cm (width) x 40 cm (height) (Maze Engineers) and allowed to explore the chamber for 5 min freely (750 lux throughout the chamber). The center region was designated as 20 × 20 cm^2^. The EthoVision XT video tracking system (Noldus) was used to record the session and analyze the behavior, movement, and activity of animals. We wiped the chamber with 95% ethanol before subsequent tests to remove any scent clues [25]. The total distance traveled and time spent in the center and outer zones of the chamber were measured.

#### Elevated plus maze test

Mice were placed in the central area of the maze consisting of four arms (two open and two closed arms) in the shape of a plus sign (35cm arm in length, 5cm arm in width, 20cm wall in height) elevated 60 cm from the ground (Maze Engineers). Mice were placed in the center region and explored the maze for 5 min freely (750 lux throughout the chamber). The session was recorded, and the number of arm entries and the amount of time spent in the open and closed arms were analyzed with the EthoVision XT video tracking system.

#### Light-dark box test

The box consists of two chambers (one light, 25cm in length x 40cm in depth and one dark, 17.5cm in length x 40cm in depth, Maze Engineers). Mice were placed in the center of light chamber and allowed to explore the chambers for 5 min freely (750 lux throughout the chamber). Time spent in each compartment and crossings from one compartment to the other were measured with the video tracking system.

#### Conditioned place preference test

The experiment involved four phases: habituation phase, pre-conditioning test, conditioning phase, post-conditioning test. Five days of free access to all environments allows the mice to habituate to the apparatus having two compartments (20cm in width x 18cm in depth x 30cm in height) with a corridor (20cm in width x 7cm in depth x 30cm in height, Maze Engineers). The last day of habituation trial data was used as pre-conditioning test data. For conditioning phase, the compartments were divided into compartments with or without food and mice introduced into one of two compartments, with restricted access for 30 min. After four days of conditioning phase, preference tests were performed. Mice were introduced into the center compartment with free access to all compartments for 15 min without food. The result of the preference test was calculated using time spent in each compartment during the pre-conditioning and post-conditioning test. The CPP score was determined by subtracting the time spent in the food-paired chamber during habituation from the time spent in the same chamber on the test day.

### Measurement of plasma corticosterone and norepinephrine levels

Blood samples were collected from the retroorbital plexus using heparinized capillary tubes (VWR international, LLC). Whole blood samples were centrifuged at 2,000 X g for 10 min at 4°C, and the plasma was separated and stored at –20 °C until use. Plasma corticosterone and norepinephrine levels were determined using two-site sandwich ELISA kits (corticosterone: Cayman, 501320; norepinephrine: MyBioSource, MBS2600834) following the manufacturer’s protocols.

### Statistics

All statistical results were presented as mean±SEM. Statistical analyses were performed using Graphpad Prism 10.0. Two-tailed *t*-tests were used to calculate p values of pair-wise comparisons. Time course comparisons between groups were analyzed using a two-way repeated-measures (RM) ANOVA with Sidak’s correction for multiple comparisons. Data were considered significantly different when the probability value is less than 0.05.

### Results

#### Deletion of the *Magel2* gene in ARC^Pomc^ neurons that innervate the MeA

We sought to determine whether knockdown of *Magel2* expression in ARC^Pomc^ neurons that innervate the MeA causes an increase in anxiety-like behavior. To achieve this, we used a retrograde adeno-associated virus (AAVrg) containing the Cre recombinase under the control of neuronal *Pomc* enhancers (nPEs) to genetically eliminate the *Magel2* gene, as we have done previously [22; 23]. We first validated our experimental approaches by injecting AAVrg-nPEs-Cre into the MeA of Rosa26-floxed-stop CRISPR-associated protein 9 (Cas9)-enhanced green fluorescent protein (eGFP) knock-in mice (Rosa26-Cas9^f^) (Fig. 1A). Three-four weeks post viral injections, double immunostaining with anti-GFP and anti-POMC antibodies revealed that a subset of ARC^Pomc^ neurons co-expressed GFP (Fig. 1B). This finding indicates that these ARC^Pomc^ neurons are those that project to the MeA, further confirming our previous studies [17; 18]. In this experimental setup, we injected AAVrg-DIO-tdTomato-U6-*Magel2* sgRNA along with AAVrg-nPEs-Cre into the MeA of Rosa26-Cas9^f^ mice to specifically knock down the expression of the *Magel2* gene in ARC^Pomc^ neurons that project to the MeA (Fig. 1 C-E). Quantification of MAGEL2-positive cells within the ARC revealed a significant decrease in the number of MAGEL2-positive neurons (*p*=0.01; Fig. 1E), indicating knockdown of *Magel2* in ARC^Pomc^ neurons innervating the MeA.

**Figure 1.**
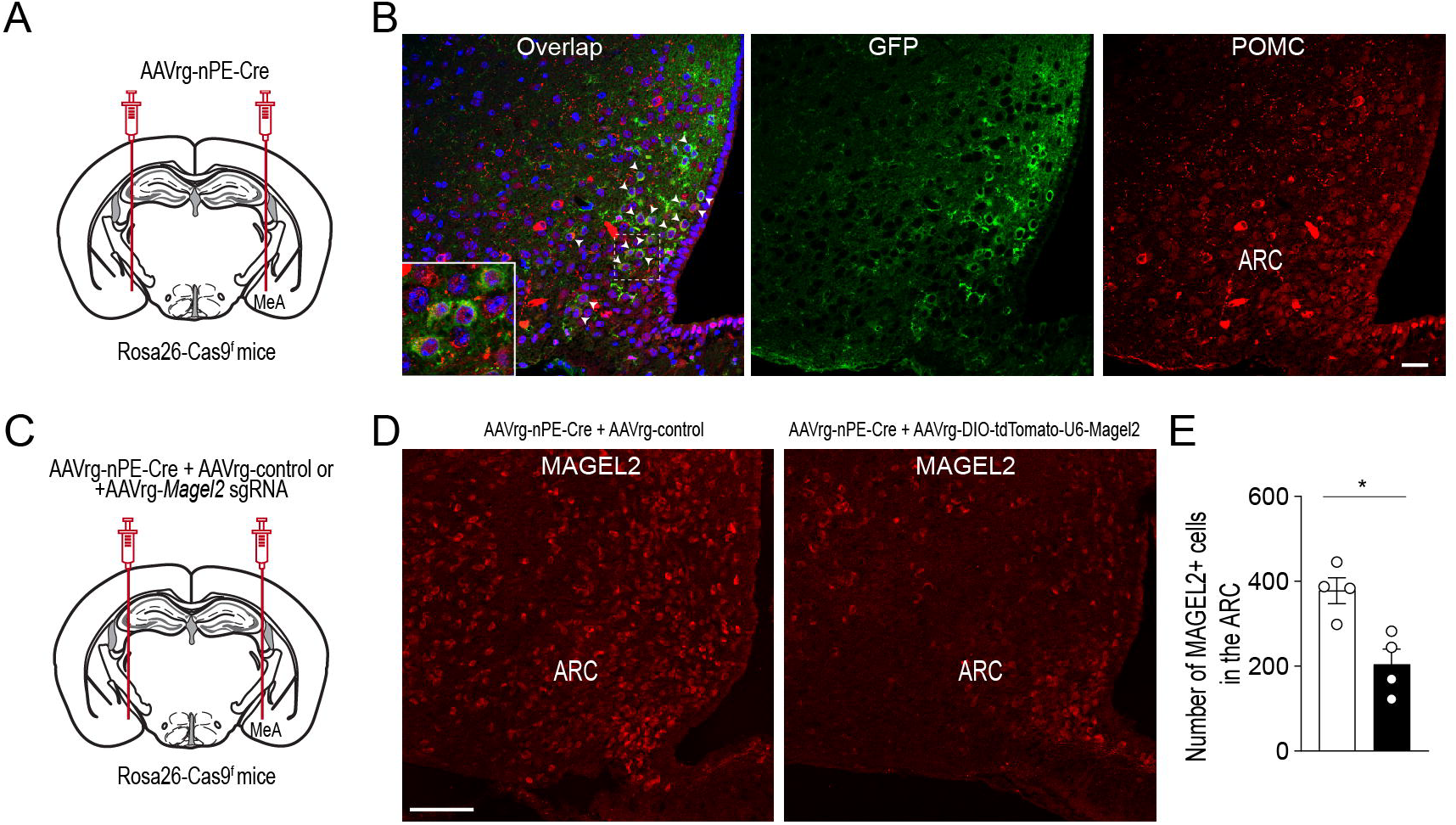
Selective deletion of *Magel2* in MeA-projecting ARC^Pomc^ neurons. (A) Schematic illustration of the experimental condition. AAVrg-nPE-Cre viruses were injected into the MeA of Rosa26-Cas9^f^ mice. (B) Images of confocal fluorescence microscopy showing co-expression of GFP and POMC in a subset of ARC^Pomc^ neurons within the ARC of Rosa26-Cas9^f^ mice injected with AAVrg-nPE-Cre. Scale bar, 30μm (C) Schematic illustration of experimental conditions. A mixture of AAVrg-nPE-Cre viruses and either AAV-control or AAVrg-*Magel2* sgRNA viruses was bilaterally injected into the MeA of Rosa26-Cas9^f^ mice. (D and E) Images of confocal fluorescence microscopy showing the expression of MAGEL2-positive neurons in Rosa26-Cas9^f^ mice injected with AAVrg-nPE-Cre and AAVrg-control or AAVrg-Magel2 sgRNA into the MeA (D, scale bar, 100 μm). Quantitative analysis of MAGEL2-positive neurons in the ARC (n = 4 mice per group) revealed a significant decrease in the number of MAGEL2-positive cells. Unpaired t-test, ^*^ *p*=0.01. Data are presented as mean values ± SEM.

#### Effect of the loss of function of the Megale2 gene in MeA-innervating ARC^Pomc^ neurons on spatial learning associated with a food reward

Male mice lacking the *Magel2* gene showed reduced learning capabilities, especially in spatial learning [26]. We performed a conditioned place preference (CPP) test, which depends on the animal’s capacity to learn and remember spatial associations between environmental cues and the rewarding effects of the paired stimulus. During the conditioning phase, Emmental cheese, as a palatable food, was associated with spatial information. On the day of testing, the food was removed, and the duration spent in the compartment previously paired with the food was measured (Fig. 2A). Results revealed that male mice lacking the *Magel2* gene in MeA-innervating ARC^Pomc^ neurons spent significantly less time in the compartment paired with food compared to the controls (*p*=0.007; Fig. 2B), while no significant difference was observed in the total distance traveled between the groups (Fig. 2B). In contrast to male mice, we found no significant differences between the female control and experimental groups (Fig. 2C and D). In other words, the time spent in the food-paired compartment was similar across these groups. Hence, these results suggest that the *Magel2* gene in ARC^Pomc^ neurons may play a role in short-term memory and spatial learning that was associated with food reward in male mice.

**Figure 2.**
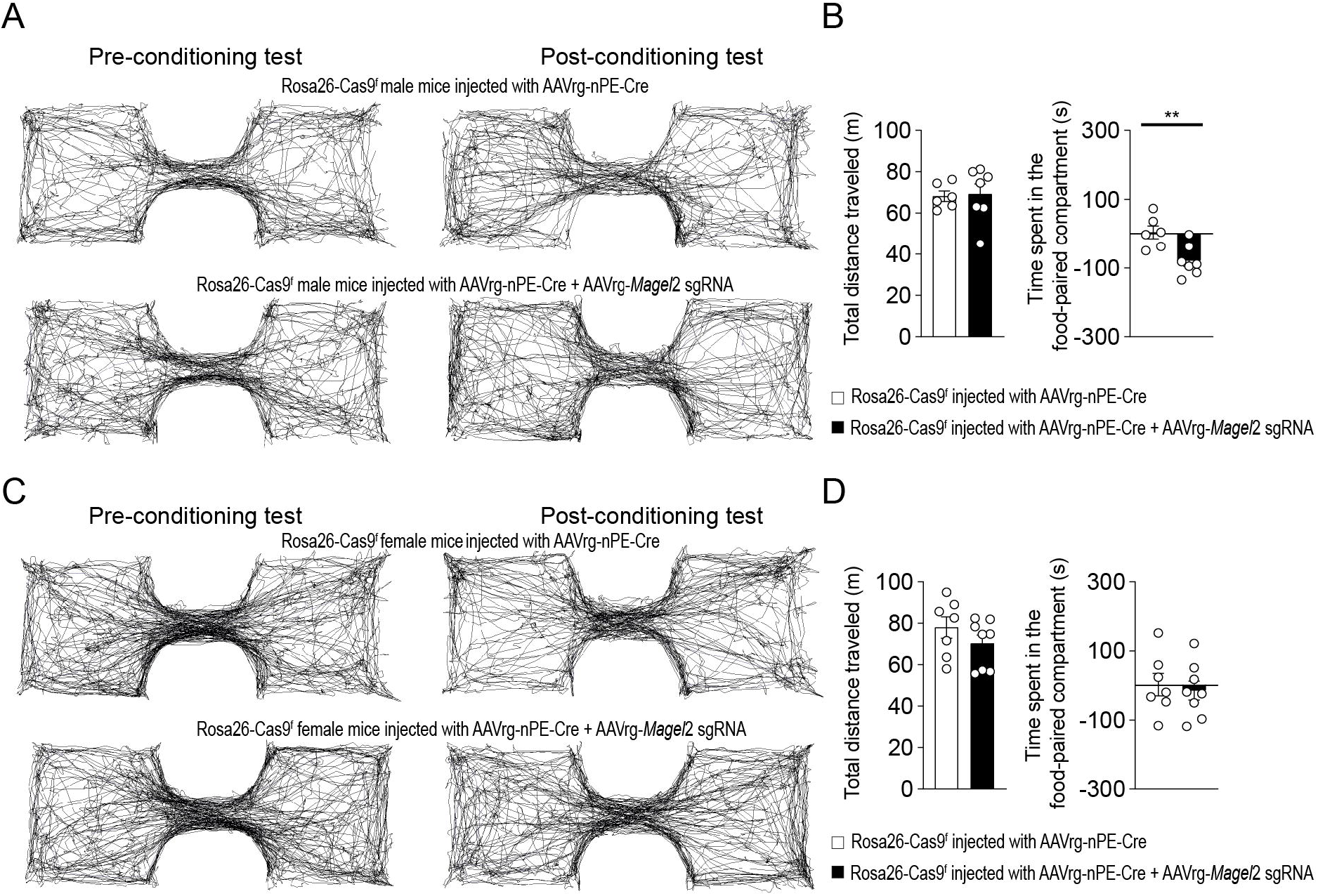
Knockdown of *Magel2* in ARC^Pomc^ neurons causes an impairment in spatial learning associated with a food reward in male mice. (A) Representative example of the travel paths of the control (top) and the experimental (bottom) groups during the CPP test in male mice. The experimental mice showed no preference for the food-paired chamber. (B) Graphs showing the total distance (left) on the test day and the CPP score between the control and experimental groups (n = 6 mice vs. 7 mice, unpaired t-test, ^**^ *p*=0.007). (C) Representative example of the travel paths of the control (top) and the experimental (bottom) groups during the CPP test in female mice. (D) Graphs showing the total distance and the time spent in the food-paired compartment in the conditioned place preference test between the female groups (n = 7 mice vs. 8 mice). No differences in the total distance (left) on the test day and the CPP score between the control and experimental groups were observed.

#### Impact of acute stress on anxiety-like behavior in mice lacking *Magel2* in MeA-innervating ARC^Pomc^ neurons

We then explored the impact of *Magel2* gene deletion in MeA-innervating ARC^Pomc^ neurons on anxiety-like behavior as anxiety is very common in individuals with PWS [7; 27]. We first examined if the loss of function of the *Magel2* gene in these neurons influences body weight and found that there were no significant differences in body weight between the groups (Fig. 3A and B). We then performed a series of experiments in male mice. Initially, we conducted an open field test, a common method for evaluating anxiety-like behavior [25; 28]. Behavioral elements of anxiety include decreased total locomotor activity, lower distance traveled and lower percentage of time spent in the center region, and higher percentage of time spent in the periphery zone. We observed no significant differences in the total distance traveled or the duration spent in the center zone between the control and experimental groups (Fig. 3C and D), which is consistent with the prior findings with *Magel2* knockout (ko) mice [26; 29; 30]. Furthermore, we conducted a light/dark (LD) box test to evaluate changes in the willingness to explore the illuminated, unprotected area, as well as an elevated plus maze (EPM) test to investigate the inherent tendency of mice to explore novel environments by offering a choice between open, unprotected maze arms and enclosed, protected arms [31]. Mice deficient in the *Magel2* gene within MeA-projecting ARC^Pomc^ neurons did not demonstrate any differences in these assessments compared to the control group in LD test (Fig. 3E and F) and EPM (Fig. 3G and H) as described in *Magel2* ko mice [30]. These findings suggest no significant differences in anxiety-like behavior between the two male groups, which is not consistent with the commonly recognized anxiety associated with PWS [27].

**Figure 3.**
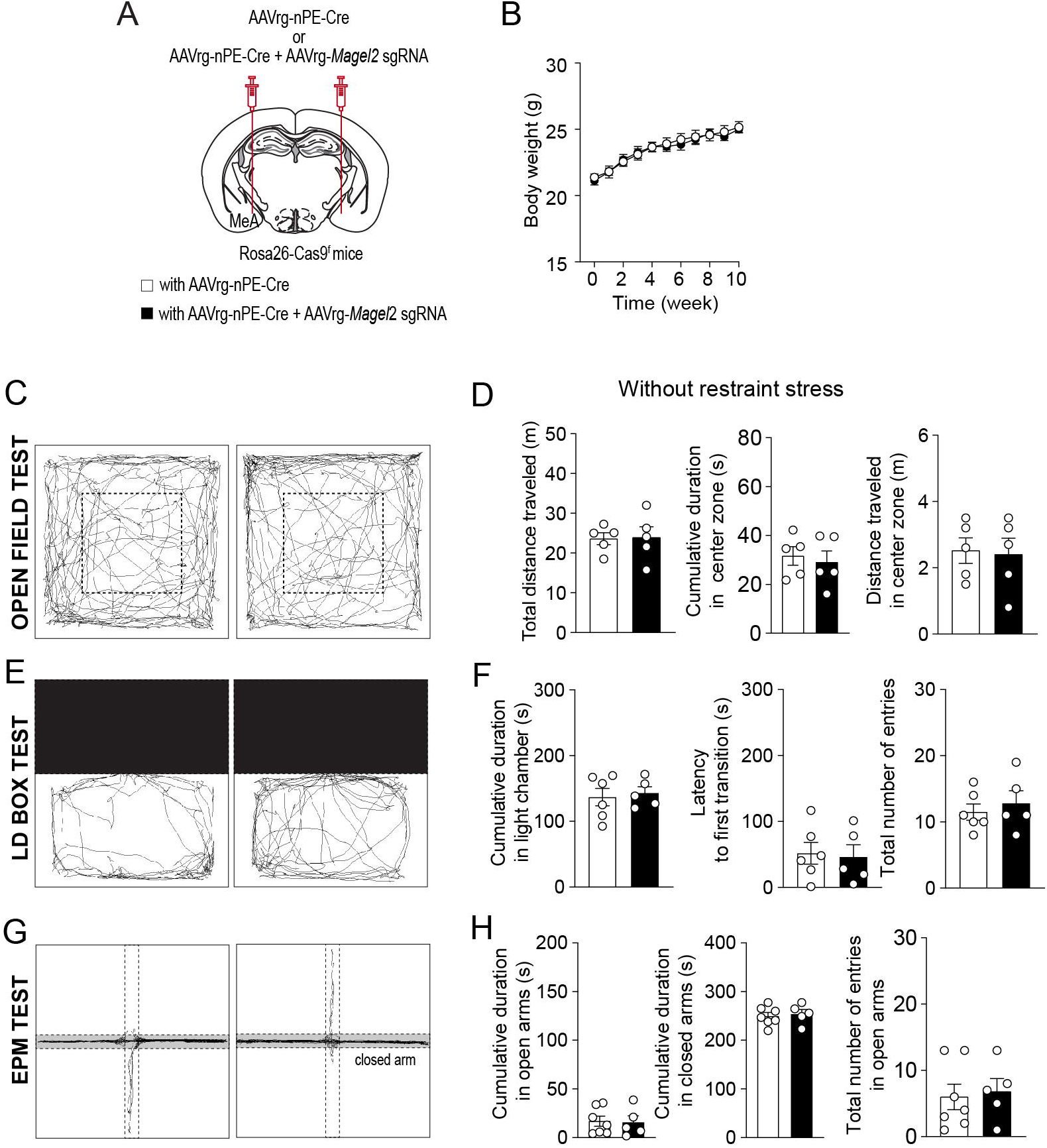
Loss of function of the *Magel2* gene in ARC^Pomc^ neurons does not alter anxiety-like behavior in male mice. (A) A schematic illustration of the experimental condition. (B) Graph showing no significant difference in body weight between the control (n= 6 mice) and experimental (n= 7 mice) groups following viral injections. Body weight was measured weekly at 9 a.m. (C and D) Representative example of the travel paths of the control (left) and the experimental (right) groups (C). Graphs showing the total distance traveled, the time spent and the distance traveled in the inner area during the open field test (n = 5 mice vs. 5 mice). (E and F) Example of the travel paths of the control (left) and the experimental (right) groups (E). Graphs showing no significant differences in the time spent in the light chamber, the latency to first transition, and the total number of entries during the light-dark box test (n = 7 mice vs. 5 mice). (G and H) Example of the travel paths of the control (left) and the experimental (right) groups (G). Graphs showing no significant differences in the cumulative duration in the open and closed arms during the elevated plus maze test (n = 7 mice vs. 5 mice).

Individuals with PWS are highly stress-sensitive [32]. We thus examined if acute stress such as restraint affects anxiety-like behavior in male mice lacking the *Magel2* gene in MeA-innervating ARC^Pomc^ neurons (Fig. 4A). In the open field test, we found that, although both the control and experimental mice exhibited similar total distance traveled (Fig. 4B and C), the mice lacking the *Magel2* gene in MeA-innervating ARC^Pomc^ neurons spent significantly less time in the center zone compared to that of the control group following short-term restraint stress (*p*=0.009; Fig. 4C). Furthermore, the results of the LD box test demonstrated that the experimental mice required a significantly longer latency to make their first transition compared to the control group (*p*=0.01; Fig. 4D and E). However, we observed that acute stress had no impact on any parameters, including the duration spent in open arms and the frequency of entries into open arms during the EPM test (Fig. 4F and G). Our findings suggest that the loss of the *Magel2* gene may influence the impact of stress on anxiety-like behavior in male mice. This observed difference in the effect of stress on anxiety-like behavior could be attributed to elevated levels of stress hormones including corticosterone and norepinephrine. However, our measurements of these hormones in plasma revealed no significant differences in their levels (Fig. 4H).

**Figure 4.**
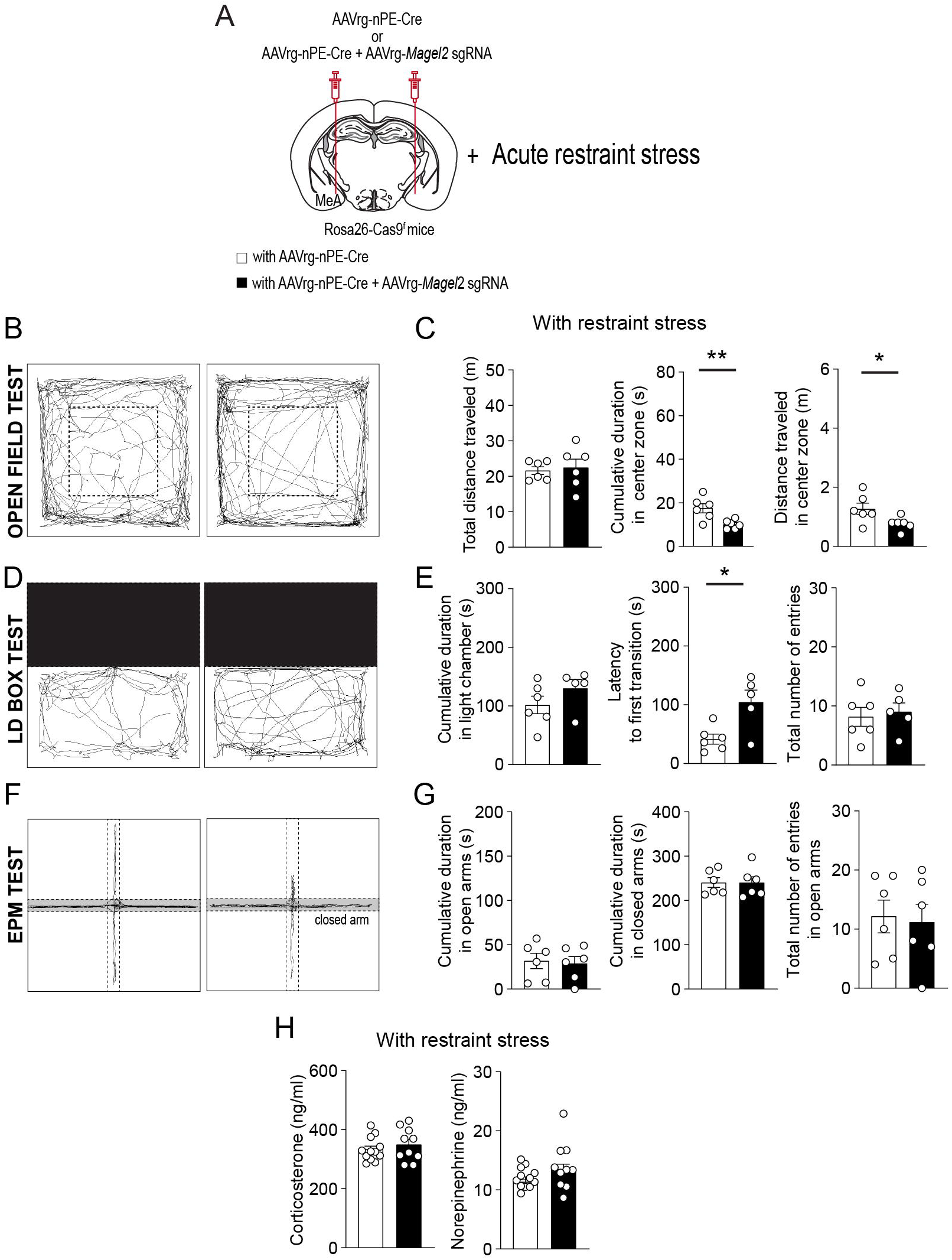
Loss of function of the *Magel2* gene in ARC^Pomc^ neurons alters the effect of stress on anxiety-like behavior in male mice. (A) A schematic illustration of the experimental condition. Mice were restrained for 30 min (B and C) Representative example of the travel paths of the control (left) and the experimental (right) groups (B). Graphs showing the total distance traveled, the time spent, and the distance traveled in the inner area during the open field test following exposure to restraint stress for 30 min (n = 6 mice vs. 6 mice, unpaired *t*-test, ^**^ *p*=0.009; ^*^*p*=0.04). (D and E) Representative example of the travel paths of the control (left) and the experimental (right) groups (D). Graphs showing the time spent in the light zone, the latency to first transition, and the total number of entries during the light-dark box test (n = 6 mice vs. 5 mice, unpaired t-test, ^*^ *p*=0.01). There was a significant difference in the latency to first transition between the groups. (F and G) Representative example of the travel paths of the control (left) and the experimental (right) groups (F). Graphs showing the cumulative duration in the open and closed arms during the elevated plus maze test with restraint stress (n = 6 vs. 6 mice). (H) Graphs showing plasma corticosterone (left) and norepinephrine (right) levels in the control (n=12 mice) and experimental (n= 10 mice) groups. No significant differences were observed between the groups. Data are presented as mean values ± SEM.

We also conducted same behavioral assessments on female mice. Similar to male mice, female mice lacking the *Magel2* gene in ARC^Pomc^ neurons innervating the MeA exhibited no difference in body weight compared to the control mice (Fig. 5A and B). Consistent with findings in *Magel2* ko female mice [26], the results from the open field test, LD box test, and EPM test in the absence of acute restraint stress exhibited no significant differences between the control and experimental mice (Fig. 3C-H). Namely, both groups displayed comparable total distance traveled and time spent in the inner zone in the open field test (Fig. 5D). No significant differences were observed in the duration spent in the light chamber or in the latency to first transition during the LD box test (Fig. 5E and F). Similarly, the EPM test showed no differences in the duration spent in the open arms or the frequency of entries into the open arms (Fig. 5G and H). In addition, we also found no significant difference in the open field test (Fig. 6A, B, and C), LD box test (Fig. 6D and E), and EPM test (Fig. 6F and G) between the two female groups following acute exposure to stress. Measurement of plasma corticosterone and norepinephrine levels showed no significant difference between the groups (Fig. 6H).

**Figure 5.**
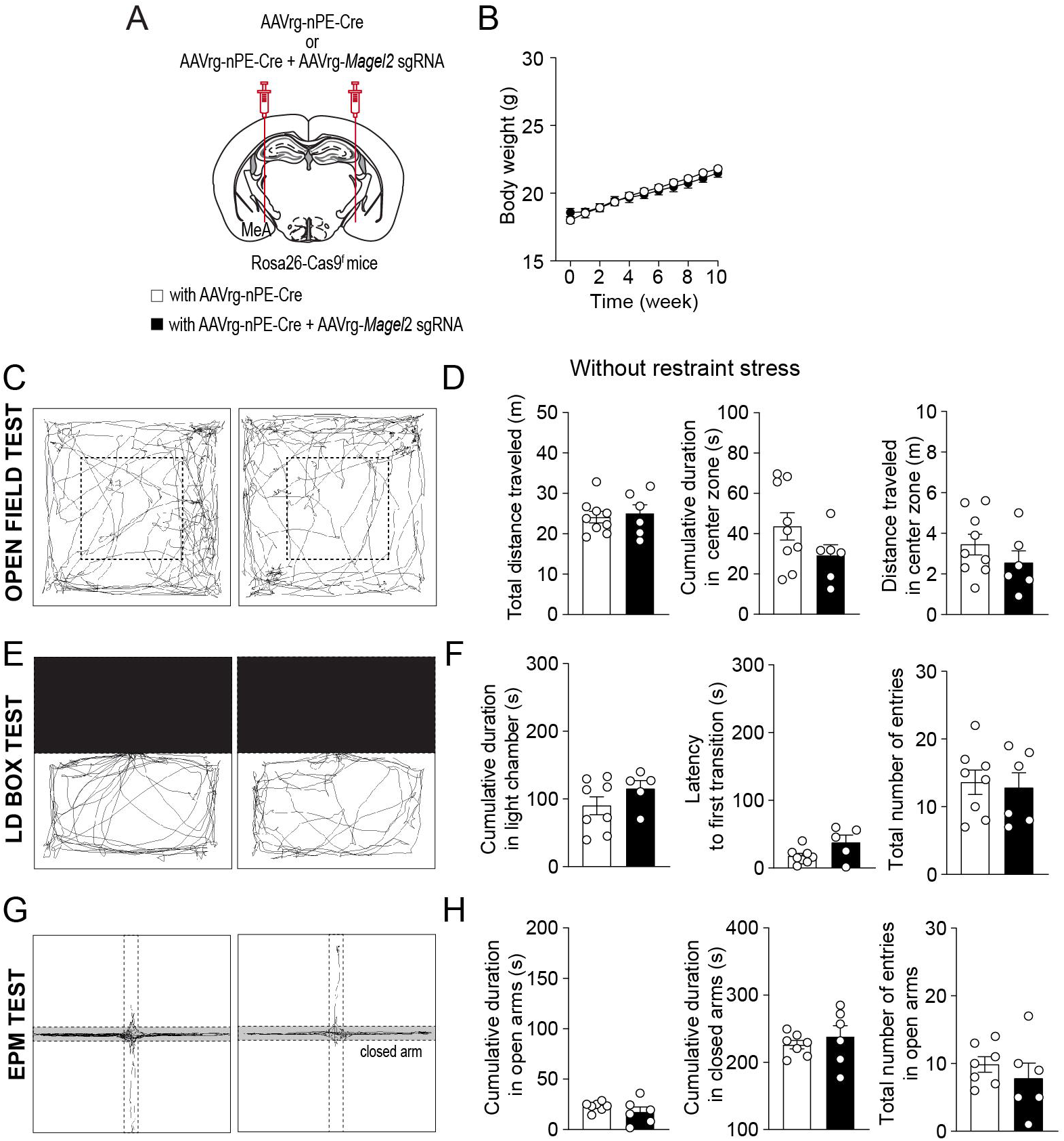
Deleting *Magel2* in ARC^POMC^ neurons does not affect anxiety-like behavior in female mice. (A) A schematic illustration of the experimental condition. (B) Graph showing no difference in body weight between the control (n= 7 mice) and experimental (n= 8 mice) groups following viral injections. (C and D) Representative example of the travel paths of the control (left) and the experimental (right) groups (C). Graphs showing no significant differences in the total distance traveled, the time spent and the distance traveled in the inner area during the open field test (n= 9 mice vs. 6 mice). (E and F) Example of the travel paths of the control (left) and the experimental (right) groups (E). Graphs showing the time spent in the light chamber, the latency to transition, and the total number of entries during the light-dark box test (n= 8 mice vs. 5 mice). (G and H) Graphs showing the cumulative duration in the open and closed arms during the elevated plus maze test (n = n= 7 mice vs. 6 mice). No significant differences were observed between the groups.

**Figure 6.**
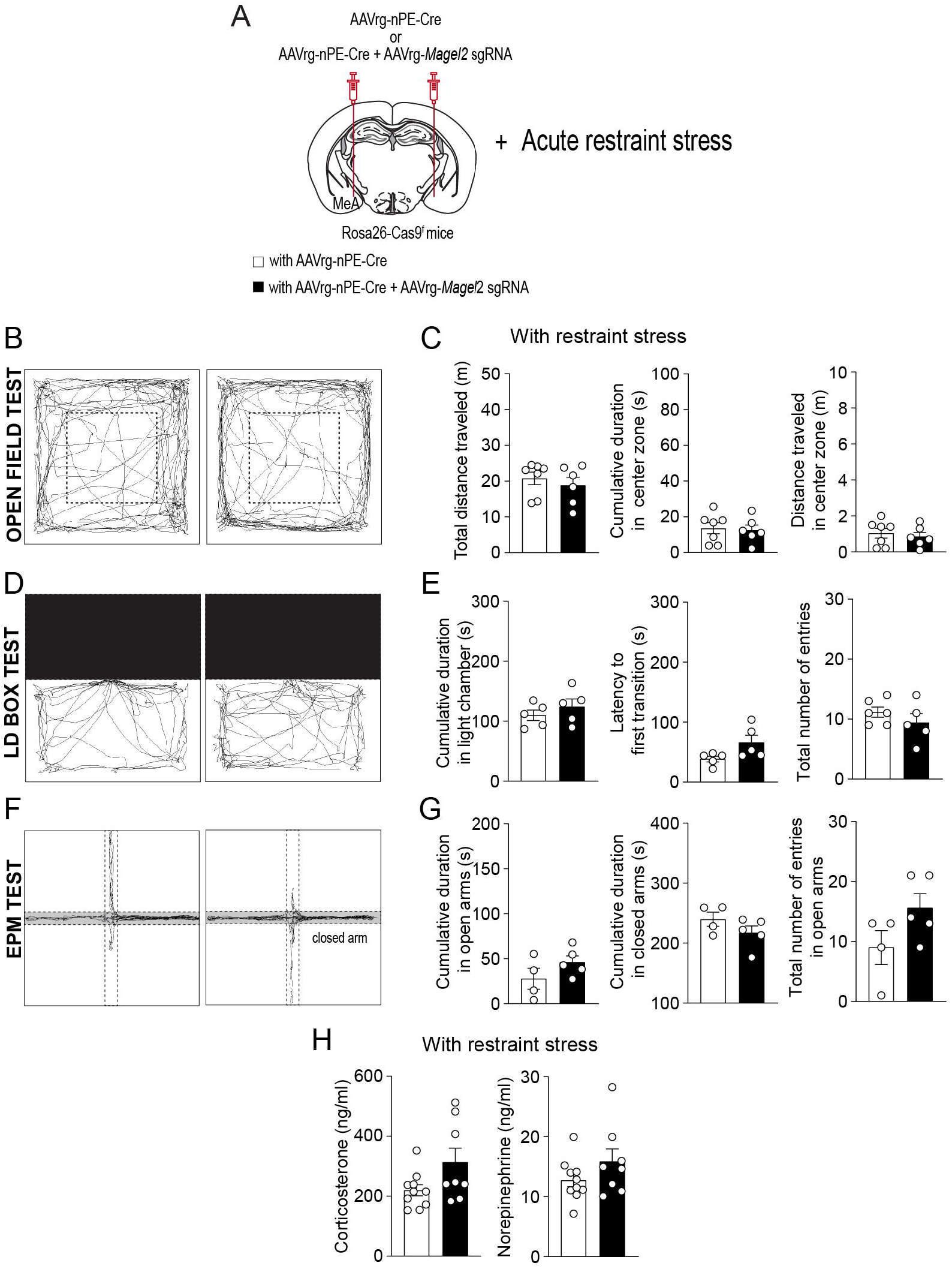
Deleting *Magel2* in ARC^POMC^ neurons does not influence the impact of stress on anxiety-like behavior in female mice. (A) A schematic illustration of the experimental condition. Mice were restrained for 30 min (B and C) Representative example of the travel paths of the control (left) and the experimental (right) groups (B). Graphs showing the total distance traveled, the time spent, and the distance traveled in the inner area during the open field test following exposure to restraint stress for 30 min (n= 7 mice vs. 6 mice). (D and E) Representative example of the travel paths of the control (left) and the experimental (right) groups (D). Graphs showing the time spent in the light zone, the latency to first transition, and the total number of entries during the light-dark box test following exposure to restraint stress (n= 5 mice vs. 5 mice). (F and G) Representative example of the travel paths of the control (left) and the experimental (right) groups (F). Graphs showing the cumulative duration in the open and closed arms during the elevated plus maze test with restraint stress (n= 4 mice vs. 5 mice). (H) Graphs showing plasma corticosterone (left) and norepinephrine (right) levels in the control (n=10 mice) and experimental (n= 8 mice) groups. Data are presented as mean values ± SEM.

## Discussion

PWS is a complex neurodevelopmental disorder characterized by a range of behavioral changes, including not only hyperphagia, but also anxiety, temper outbursts, and obsessive-compulsive behavior [7; 27]. The cellular mechanisms underlying the behavioral changes in PWS has been studied in animal models such as *Magel*2 ko mice [26; 29; 30]. In this study, we specifically deleted the *Magel2* gene in ARC^Pomc^ neurons that innervate the MeA, given that the MeA is a critical structure involved in emotional behaviors, particularly anxiety-like behavior [33]. We found that both male and female mice lacking the *Magel2* gene in MeA-innervating ARC^Pomc^ neurons did not display any significant alterations in the open field maze, LD box, and EPM assessments in the absence of restraint stress. In contrast, the male mutant mice exhibited increased anxiety-like behaviors such as a reduced time spent in the center zone and a latency to first transition, compared to their control littermates following exposure to restraint stress. Moreover, the loss of function of the *Magel2* gene in these specific ARC^Pomc^ neurons resulted in decreased spatial learning that was associated with a food reward in male mice. Contrary to the male mutant mice, the female mutant mice did not show any behavioral changes with and without exposure to restraint stress. Given that individuals with PWS are highly sensitive to stress, which can lead to behavioral changes [32], our present study may offer a novel perspective on the role of the *Magel2* gene, among other imprinted genes associated with PWS, in regulating the impact of stress on anxiety-related behaviors within this population.

Hyperphagia is a primary characteristic of PWS, resulting in excessive weight gain and obesity. Intriguingly, the loss of function of the *Magel2* gene did not replicate this characteristic feature in two distinct *Magel2* ko mouse models, namely the LacZ knock-in allele and *Magel2* deletion [10; 19; 30; 34]. Although the male *Magel2* ko mice with the LacZ knock-in allele showed a slight increase in body weight on a standard chow diet [19], the *Magel2* ko mice did not develop DIO compared to their control littermates when fed a high-fat diet (HFD). Instead the *Magel2* ko mice demonstrated a reduction in body weight relative to the control mice during HFD feeding [19]. *Magel2* ko mice with *Magel2* deletion also showed normal weight, temperature, motor abilities, and pain sensitivity [26]. These two distinct ko mice displayed a reduction in food intake compared to the controls [10; 19], which seems to be partially attributed to changes in circadian output [10]. In our prior study [18], deleting the *Magel2* gene in ARC^Pomc^ neurons that project to the MeA caused a reduction in body weight in both male and female mice compared to the control littermates on HFD feeding. This was associated with increased locomotor activity particularly in the dark phase. Additionally, these mutant mice did not exhibit any differences in food intake, glucose tolerance, and insulin sensitivity [18]. Hence, these prior studies suggest that the *Magel2* gene, among the imprinted genes associated with PWS, might not be crucial in the development of hyperphagia and obesity.

Despite the *Magel2* gene not affecting hyperphagia, the *Magel2* ko mice displayed altered behavioral phenotypes, such as a lack of preference for social novelty, a deficit in social interaction and recognition, and a reduced learning ability in a sex-dependent manner [26; 29; 30]. Interestingly, although anxiety is the most prevalent behavioral change in PWS [27], *Magel2* ko mice did not exhibit anxiety-related behavior in the open field maze [26; 29] and elevated plus maze assessments [30]. Contrary to these findings, it has been also demonstrated that both male and female *Magel2* ko mice spent more time in the open arms of the elevated plus maze compared to the control group, indicating changes in their exploratory behavior [29]. In this study, we found that deleting the *Magel2* gene in MeA-innervating ARC^Pomc^ neurons also did not affect anxiety-like behavior in both male and female mice in the absence of acute stress exposure, aligning with prior findings [26; 30]. According to the foundation for Prader-Willi research [32], individuals with PWS are highly susceptible to stress. Following short-term exposure to restraint stress, our male mutant mice exhibited reduced time spent in the central area during the open field test and demonstrated increased latency to first transition during the light-dark box test, indicating an elevation in anxiety-like behavior, whereas the female mutant mice did not display any alterations in anxiety-like behavior. In the Pavlovian fear-conditioning test, *Magel2* ko mice exhibited significantly higher absolute freezing rates following the tone compared to wild-type controls, suggesting increased anxiety [30]. Hence, our current findings, along with previous ones, support the idea that the loss of function of the *Magel2* gene may contribute to an enhanced impact of stress on anxiety-like behavior.

It has been shown that acute emotional stressors such as restraint and forced swim led to increased c-fos mRNA expression mainly in ARC^Pomc^ neurons rather than another important feeding-related neurons, agouti-related peptide-expressing neurons in the ARC [20]. Both stressors resulted in a reduction of food intake and an increase in anxiety-like behavior, effects that were entirely inhibited by the administration of a melanocortin receptor type 4 (MC4R) antagonist. [20]. Furthermore, MC4R-expressing neurons in the MeA were activated by acute stress exposure and inhibiting MC4Rs locally in the MeA abolished stress-induced anxiety-like behavior [35]. We previously showed that activation of the ARC^Pomc^ −>MeA pathway reduced food intake through MC4Rs [17]. Thus, it is possible that mice lacking the *Magel2* gene in MeA-innervating ARC^Pomc^ neurons may be more susceptible to stress compared to the control groups.

Intriguingly, male mice deficient in the *Magel2* gene also demonstrated a significant impairment in object recognition memory and a decrease in spatial learning abilities, whereas female mutant mice did not show these deficits [26]. In the CPP test, the animal utilizes spatial cues, such as walls with black stripes in our experimental setup, to establish a memory of the compartment associated with palatable food. Subsequently, the animal demonstrates a preference for this location based on the learned association. Our results revealed that male, but not female, mice lacking the *Magel2* gene in MeA-innervating ARC^Pomc^ neurons spent considerably less time in the compartment associated with food compared to the control mice, aligning with the previous findings [26]. The findings suggest that the *Magel2* gene, particularly within the ARC^Pomc^ neurons, may play a crucial role in short-term memory or spatial learning abilities associated with positive rewards.

The FDA has recently approved VYKAT™ XR (diazoxide choline, Soleno Therapeutics) as a treatment for hyperphagia in PWS. As MAGEL2 in ARC^Pomc^ neurons appears to be a key player in enhancing the effect of stress on anxiety-like behavior, understanding the cellular mechanisms of how MAGEL2 in ARC^POMC^ neurons innervating the MeA controls mental disorders will open up new therapeutic strategies for the treatment of mental illness in individuals with PWS.

## General

We thank Drs. Patrick Potts and Klementina Fon Tacer for providing us with the MAGEL2 antibody.

## Author contributions

SB carried out viral injection, immunostaining, behavioral tests, and ELISA assay. Y.-H. designed research, performed viral injection immunocytochemistry, analyzed the data, and wrote the manuscript.

